# MiMeNet: Exploring Microbiome-Metabolome Relationships using Neural Networks

**DOI:** 10.1101/2020.12.15.422873

**Authors:** Derek Reiman, Brian T. Layden, Yang Dai

## Abstract

The advance in microbiome and metabolome studies has generated rich omics data revealing the involvement of the microbial community in host disease pathogenesis through interactions with their host at a metabolic level. However, the computational tools to uncover these relationships are just emerging. Here, we present MiMeNet, a neural network framework for modeling microbe-metabolite relationships. Using ten iterations of 10-fold cross-validation on three paired microbiome-metabolome datasets, we show that MiMeNet more accurately predicts metabolite abundances (mean Spearman correlation coefficients increase from 0.108 to 0.309, 0.276 to 0.457, and -0.272 to 0.264) and identifies more well-predicted metabolites (increase in the number of well-predicted metabolites from 198 to 366, 104 to 143, and 4 to 29) compared to state-of-art linear models for individual metabolite predictions. Additionally, we demonstrate that MiMeNet can group microbes and metabolites with similar interaction patterns and functions to illuminate the underlying structure of the microbe-metabolite interaction network, which could potentially shed light on uncharacterized metabolites through “Guilt by Association”. Our results demonstrated that MiMeNet is a powerful tool to provide insights into the causes of metabolic dysregulation in disease, facilitating future hypothesis generation at the interface of the microbiome and metabolomics.

## Introduction

The microbiome has been shown to impact both host development, normal metabolic processes, as well as the pathogenesis of various diseases [1-3]. Of particular interest is the microbiome of the human gut, which has been linked to diseases such as inflammatory bowel disease (IBD), obesity, and diabetes mellitus [4-7]. While previous studies have uncovered various microbe-disease associations, recent work also revealed the central role of bacterial metabolites and their impact on host health [8-13]. The microbiome-metabolome crosstalk is pervasive. For example, strong associations between microbes and metabolites were found in the gut and blood metabolomic profiles [14] and the gut of patients with IBD [15]. The abundance of metabolic pathways is relatively consistent despite considerable variability in taxonomic composition among individuals [16-19]. Thus, the identification of mechanisms of microbiome-metabolome interactions by modeling community metabolic activity is essential for understanding how the microbiome affects the host’s health and for the development of precise therapies for the prevention or management of chronic diseases [20, 21].

Early methods for modeling have focused on mapping metagenomic features to metabolomic features due to the unavailability of metabolomic profiles. One set of methods referred to as Predictive Reactive Metabolic Turnover (PRMT) uses known enzymatic reactions from genome annotations and existing metabolic pathways to calculate the relative production and consumption of metabolites using metagenomic gene abundance profiles [22, 23]. Other methods have focused on constraint-based stoichiometric modeling using flux balance analysis to learn the flux rate of metabolites in the community [24, 25]. A major limitation of these approaches is their reliance on annotated references. Missing or incorrect annotations can lead to poor performing models. Additionally, the reliance on annotations makes it difficult to identify novel mechanisms.

More recently, several machine learning models have been developed to map metagenomic features to metabolites as both data have become increasingly available. One method, MelonnPan, uses the Elastic Net linear regression to model the relative abundance of each metabolite using metagenomic taxonomic or functional features [26]. The primary emphasis of MelonnPan is predictive modeling of metabolites so that the learned models can be used for the prediction of metabolomes in similar studies where only microbiome is available. MelonnPan displayed promising performance, however, it models each metabolite individually, limiting its ability to use shared information across metabolomic features to boost prediction performance. Another method, mmvec, uses neural networks to estimates the conditional probability that a metabolite is present given the presence of a specific microbial sequence [27]. mmvec focuses on learning embeddings of microbial sequences and metabolites to capture microbe-metabolite interactions, thus, it cannot predict the entire metabolomic profile. Another recent model based on neural network encoder-decoder (NED) has proposed constraints of sparsity and non-negative weights for mapping microbiomes to metabolomes [28, 29]. The use of non-negative weights in NED imposes a stringent constraint on the model, which may limit the learning capacity. Consequently, these existing models have not fully explored the microbiome-metabolome interactions that can be uncovered through integrative analysis.

In this work, we present MiMeNet (<u>Mi</u>crobiome-<u>Me</u>tabolome <u>Net</u>work), a multi-layer perceptron neural network (MLPNN) that models the community metabolome profile using metagenomic taxonomic or functional features obtained from a microbiome sample. The use of MLPNN allows MiMeNet to be scalable regarding the amount of metagenomic and metabolomic features in the two types of omics data. Furthermore, by learning multiple tasks simultaneously, the underlying information can be transferred to augment the learning of similar tasks [30], thus leading to more robust predictive models and hence more reliable microbiome-metabolome interactions. Here, we use three paired metagenomic-metabolomic datasets obtained from studies on IBD, cystic fibrosis, soil wetting environment as well as additional external datasets of IBD to evaluate MiMeNet’s ability to predict the metabolomic profile from metagenomic features by comparing with the existing methods. Using the IBD datasets, we further show how MiMeNet uses learned network models to generate hierarchies of metagenomic and metabolomic modules, which further reveal microbe-metabolite interactions and their associations to host IBD status. The MiMeNet package and the datasets used for evaluation are freely available at https://github.com/YDaiLab/MiMeNet.

## Results

### MiMeNet framework

MiMeNet is an integrative analysis framework for microbiome and metabolome data utilizing MLPNN, which trains models to accurately predict the metabolome based on a microbiome sample represented by the microbial taxonomic composition or microbial gene features. With learned network models, MiMeNet performs additional analysis to extract interaction scores between all microbe-metabolite pairs and organizes microbes and metabolites into modules according to the patterns of interaction scores. These modules can be used to examine the microbe-metabolite interactions between modules and assess the statistical associations of modules to host phenotypes. MiMeNet takes advantage of multi-task learning (MTL) by using the entire microbiome to model the abundance of all metabolites in a unified neural network architecture. MTL is a learning approach where information in related tasks is used to improve the generalization performance of all the tasks. This approach enables MiMeNet to train more robust models. An overview of the MiMeNet framework is shown in Fig 1.

**Figure 1.**
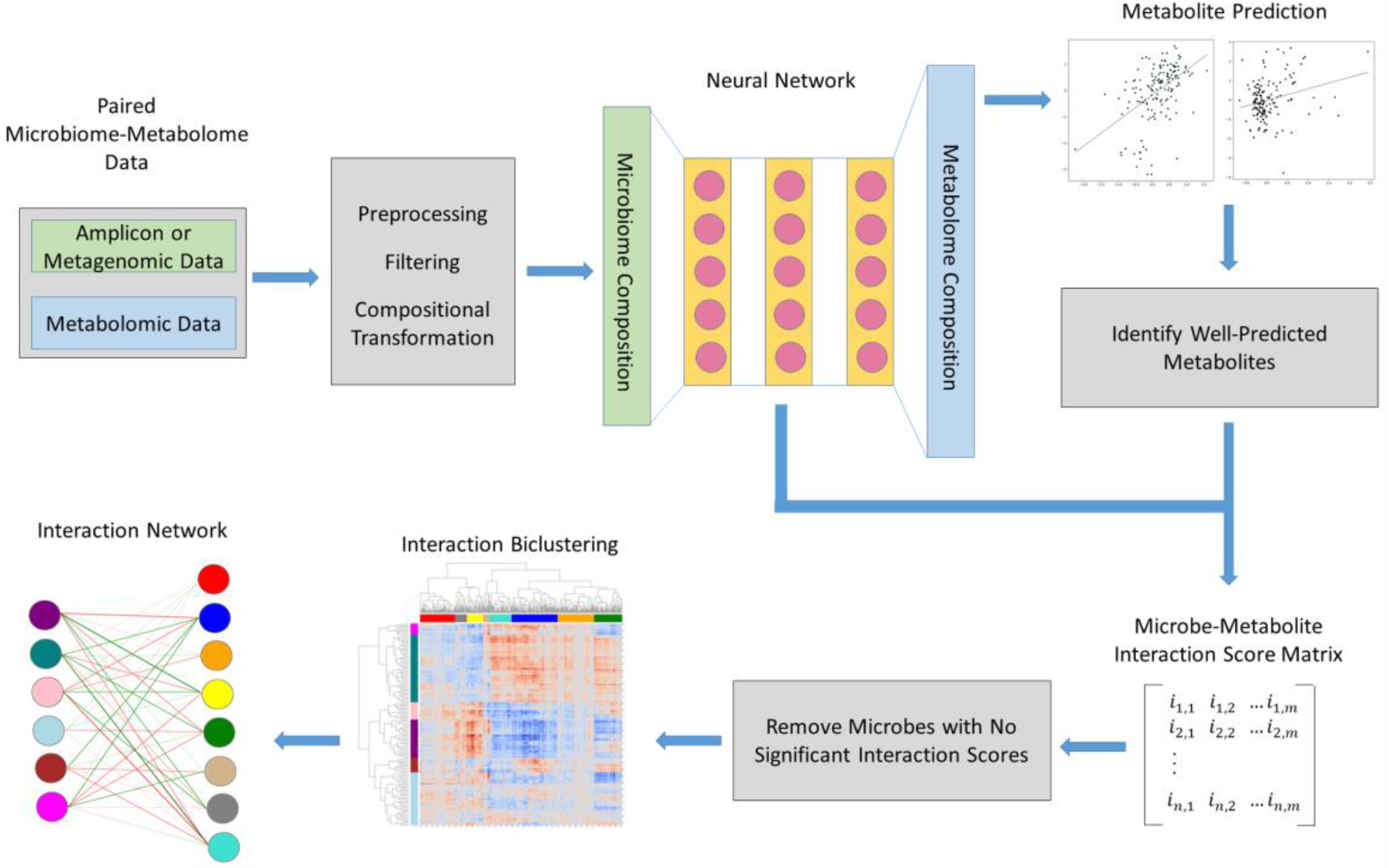
Framework of MiMeNet learning model. MiMeNet uses paired microbiome and metabolome data as input. Microbiome abundance features (green) are used to train a neural network to predict metabolite abundance features (blue). Well-predicted metabolites are identified, and the trained models are used to learn a microbe-metabolite interaction matrix. The interaction matrix is biclustered into microbe and metabolite modules which are then used to construct a module-based interaction network.

The network architecture in MiMeNet is first determined for each paired dataset by tuning the hyperparameters for the number and size of the hidden layers, the L2 regularization penalty parameter, and the dropout rate. MiMeNet then trains multiple network models using 10-fold cross-validation and the predictive performance of a model is measured by the averaged Spearman correlation coefficients (SCCs) between the predicted and the observed abundance of the metabolites across the samples in testing sets. The protocols for MLPNN model training are outlined in the Methods section. MiMeNet then generates a background distribution of SCCs through multiple iterations of shuffling the dataset and performing a cross-validated evaluation on the shuffled set. This background distribution of SCCs is then used to determine a cutoff for significantly well-predicted metabolites. Then, using the learned network weights obtained from cross-validation training, MiMeNet constructs a microbe-metabolite interaction score matrix between the microbes and well-predicted metabolites. Then MiMeNet biclusters the interaction score matrix into microbial and metabolomic modules, grouping microbes or metabolites with shared interaction patterns and constructs a module-based interaction network. Additionally, MiMeNet trains a final predictive model to be used to predict the metabolomic profile of an external set with similar microbial features (Methods).

### MiMeNet identifies significantly correlated metabolites

We used three datasets for the development and evaluation of MiMeNet (Table 1). The first dataset was taken from a published study of patients enrolled in PRISM (the Prospective Registry in IBD Study at Massachusetts General Hospital) containing 121 IBD patients and 34 controls [15]. Additionally, it includes an external test set of 43 IBD and 20 control subjects using two other cohorts. The second dataset was taken from a study that collected lung sputum samples from 172 patients with cystic fibrosis [31]. The third dataset was taken from biocrust soil water and captures microbial and metabolic activity caused by soil wetting at five-time points across four biocrust successional stages [32]. For IBD, cystic fibrosis, and soil datasets, the numbers of microbes are 201, 657, and 446, respectively; and the numbers of metabolites are 8848, 168, and 85 respectively (see Methods for details).

**Table 1.** Summary of the datasets used in evaluation. The patients of IBD have two subtypes: Crohn Disease (CD) and Ulcerative Colitis (UC).

|  | #Case | #Control | #Microbes | #Metabolites |
| --- | --- | --- | --- | --- |
| IBD (PRISM) | 68 (CD), 53 (UC) | 34 | 201 | 8848 |
| IBD (External) | 20 (CD), 23 (UC) | 20 | 201 | 8848 |
| Cystic Fibrosis | 172 | - | 657 | 168 |
| Soil | - | 19 | 446 | 85 |

Using the background distributions generated by MiMeNet (Method), the correlation cutoffs were found to be 0.136, 0.129, and 0.410 for the IBD (PRISM), cystic fibrosis, and soil datasets, respectively. Based on these cutoff values, MiMeNet identified metabolites to be well-predicted for 6857 (77.50%) of the 8848 metabolites in the IBD PRISM dataset, 143 (94.08%) of the 152 metabolites in the cystic fibrosis dataset, and 29 (34.12%) of the 85 metabolites in the soil dataset. The distributions of the background and observed correlations for the three datasets are shown in Fig 2(A-C). The soil dataset had the lowest percent of well-predicted metabolites, which could be due to the larger cutoff. We suspect that this is from the bootstrapping procedure being performed on the dataset of small size as well as the fact that the dataset is longitudinal and samples may be correlated with each other. Our evaluation shows the strong predictability of the MiMeNet trained models on data with reasonable sample sizes.

**Figure 2.**
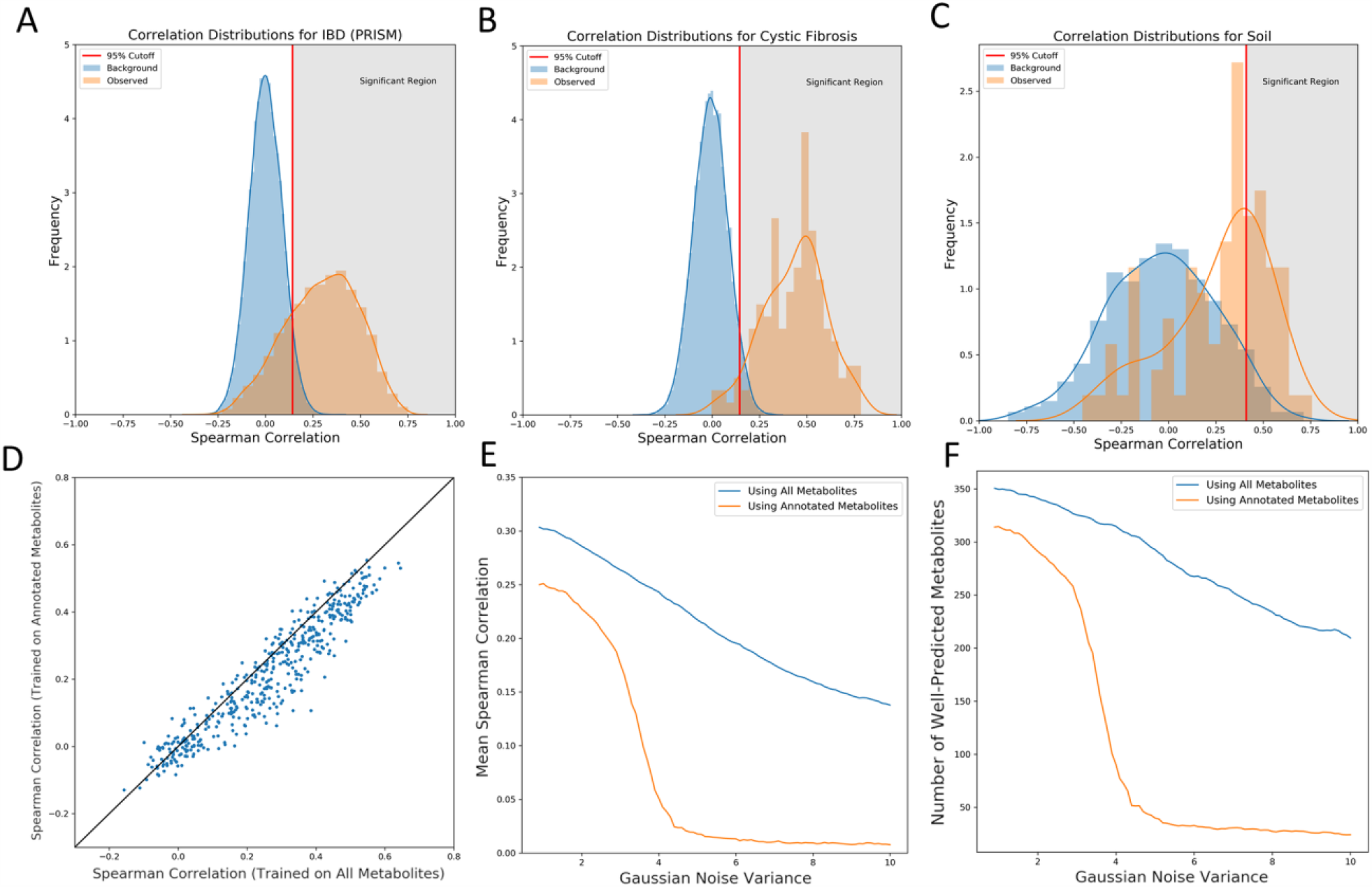
Distributions of correlations (Background and observed) and evaluation of Multi-task learning. Background (blue) and observed (orange) distributions are shown for the (A) IBD (PRISM), (B) cystic fibrosis, and (C) soil datasets. The red vertical line denotes the 95^th^ percentile of the background correlations and the gray area represents the well-predicted region using this threshold. (D) Scatter plot comparing the annotated metabolite correlations between models trained on just the annotated set and models trained on the full set of metabolites. (E) Mean correlation and (F) number of well-predicted metabolites found in models trained on the annotated set of metabolites and full set of metabolties as Gaussian noise is added to the annotated metabolite set input. All results in (D)-(F) are for prediction of the annotated metabolites.

### Multi-task learning boosted MiMeNet’s prediction performance

To evaluate if multi-task learning improves the prediction of the metabolomic profiles, we trained two separate models using 10 iterations of 10-fold cross-validation using the IBD (PRISM) dataset. The first model was trained to predict the entire set of metabolites, and the second model was trained to predict the 466 annotated set of metabolites without including the rest of the metabolites. We then compared the SCCs of the 466 metabolites from both models and observed that by training on the entire set of metabolites. The number of well-predicted metabolites for the annotated set increased from 333 to 366. Additionally, the SCCs of the annotated metabolites significantly increased from 0.259 to 0.309 when using all the metabolites to train MiMeNet (P < 10^−47^, the Wilcoxon signed-rank test). The scatter plot comparing the prediction correlation performances is shown in Fig 2D.

Next, we evaluated the robustness of MiMeNet by gradually increasing noise to the annotated set of metabolites. Specifically, using 10-fold cross-validation, we trained models using all the metabolites and using only the annotated set to predict the 466 annotated set. For each partition of the cross-validated training, we added Gaussian noise to the annotated metabolites within the training data. We observed that the two models performed similarly under small amounts of noise. However, once the noise increased to higher levels and had a variance greater than 2, the models trained only on the annotated set collapsed and could no longer predict the annotated metabolites. On the other hand, the models trained using all the metabolites were much more robust to the noise at higher levels and could predict the annotated metabolites to a much greater degree compared to those trained using only the annotated set (Fig 2(E-F)). These results show MiMeNet benefited from multi-task learning.

### MiMeNet is robust to training dataset size

To evaluate MiMeNet performance on different sizes of data for training and testing, we compared the k-fold cross-validated prediction correlations (k=10, 5, 3, and 2) using the IBD (PRISM) and cystic fibrosis datasets (the soil dataset was excluded from this analysis due to the small data size). In the IBD (PRISM), we only observed a slight decrease in performance (mean correlation coefficient decrease from 0.297 to 0.218) as the number of partitions decreased (Fig 3A). Similarly, in the cystic fibrosis dataset, the correlation dropped slightly from 0.457 to 0.410 (Fig 3B). Additionally, we evaluated performance on random subsetting of 100%, 80%, 60%, and 40% of the entire datasets. For each level of subsetting, 3 random sets of subset data were generated. Then, for each set of data, network hyper-parameters were tuned and 10 iterations of 10-fold cross-validation was performed to evaluate how reducing the number of overall samples affected the prediction correlation (Fig 3C). As the size of the dataset decreased, we observed a decrease in the IBD (PRISM) dataset from a mean correlation of 0.287 to 0.179, and a decrease in the cystic fibrosis dataset from a mean correlation of 0.443 to 0.364. Moreover, we evaluated the IBD (External) dataset for each MiMeNet model trained on the IBD (PRISM) dataset and observed a decrease in mean correlation from 0.222 to 0.162. Even though there was a decrease in overall correlations as expected, we show that MiMeNet can still predict the metabolomic profile when using smaller sets of training data.

**Figure 3.**
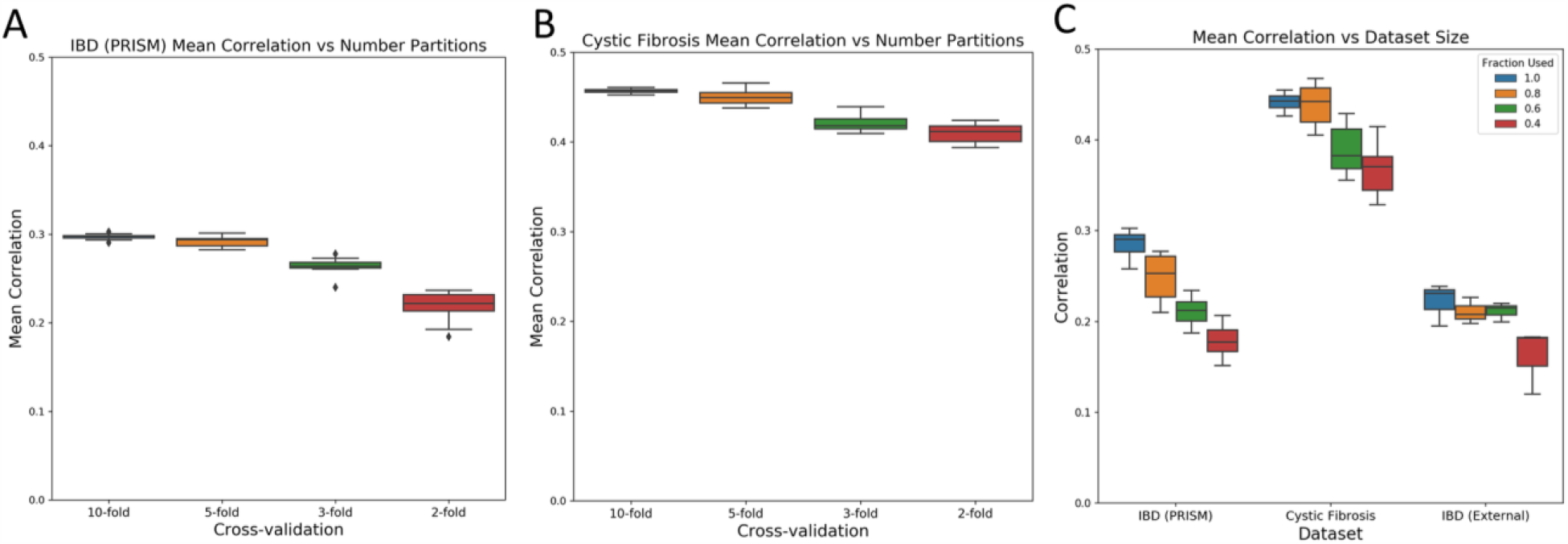
Mean correlation analysis using different amounts of training data. Correlations for 10-, 5-, 3-, and 2-fold cross-validation evaluations are shown for the (A) IBD (PRISM) and (B) cystic fibrosis datasets. (C) Subsets of the IBD (PRISM) and cystic fibrosis corresponding to 100%, 80%, 60%, and 40% of the total samples are used as an input for MiMeNet. Three random datasets for each level of subsetting were created and then mean correlation using 10 iterations of 10-fold cross-validation is calculated across the three. In addition, models trained on the complete subsets of the IBD (PRISM) data are used to evaluate the IBD (External) test set.

### MiMeNet outperforms models in MelonnPan

For benchmarking, we compared MiMeNet to MelonnPan, a recent model that uses Elastic Net linear regression to predict metabolite abundance from microbial abundance features [26]. Elastic net regression applies a linear combination of both L1 and L2 regularizations to avoid overfitting. In the case of MelonnPan, a linear model is trained for one metabolite at a time and cannot benefit from multi-task learning. In our study, MelonnPan was evaluated using the same data partitions of the 10 iterations of 10-times cross-validation for each dataset. However, in the case of the IBD (PRISM) dataset, only the annotated metabolites were predicted due to the large running time for the entire metabolite set. On the other hand, MiMeNet was trained to predict all metabolites in the IBD (PRISM) dataset. We further evaluated the performance of the models obtained from MelonnPan and MiMeNet using the IBD (External) dataset on the annotated metabolites. We observed that in each of the datasets trained using cross-validation (Fig 4(A-C)), MiMeNet has a higher correlation for prediction across all metabolites when compared to MelonnPan. In the IBD (PRISM) dataset, the mean correlation increased from 0.108 to 0.309 when evaluating the annotated metabolites. When training MiMeNet only on the annotated metabolites, we observed a similar result with an increased correlation to 0.259 (S1 Fig). In the cystic fibrosis dataset, the mean correlation increased from 0.276 to 0.457. In the soil dataset, the mean correlation from MelonnPan was -0.272 and was increased to 0.264 using MiMeNet. Moreover, when evaluating the IBD (External) dataset, the mean correlation of the annotated metabolites was increased from 0.168 to 0.275 (Fig 4D). Among the top 20 well-predicted metabolites from MiMeNet from the IBD (PRISM) dataset (Fig 4E), they are fatty acids (eicosatrienoic acid, docosapentaenoic acid, adrenic acid, and docosapentaenoate), and bile acids (chenodeoxycholate and cholate). Both of these classes of metabolites have been associated with IBD in previous studies [33-35].

**Figure 4.**
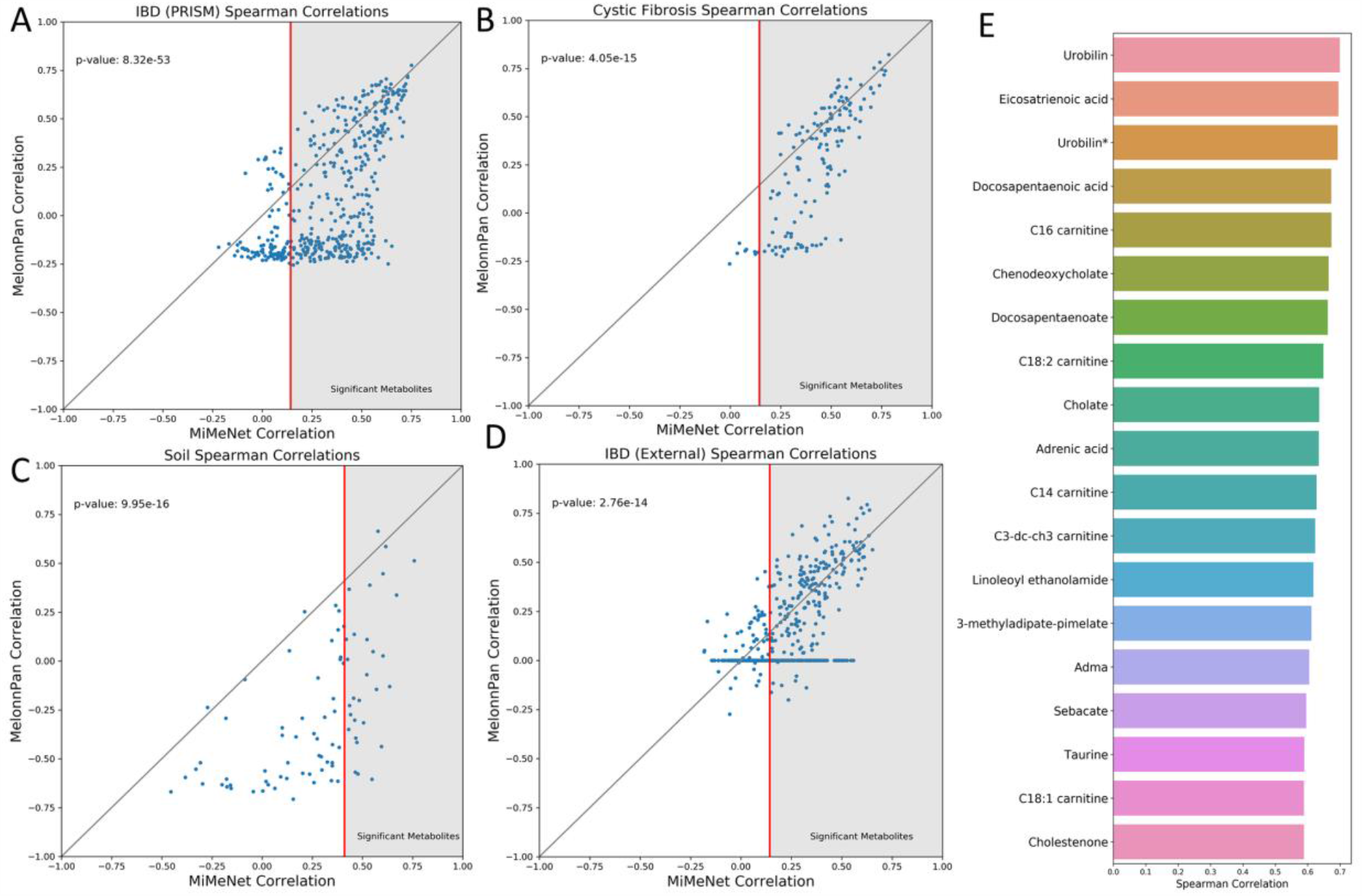
Scatter plots comparing MiMeNet to MelonnPan prediction correlations. Scatter plots showing the metabolite prediction correlations of MiMeNet against MelonnPan for (A) PRISM IBD dataset, (B) cystic fibrosis dataset, and (C) soil dataset, each trained using 10 iterations of 10-fold cross-validation. (D) In addition, 10 iterations of both models were trained on the PRISM dataset to evaluate the external IBD dataset. Each point represents the average correlation of a metabolite across 10 iterations of training. (E) The top 20 best correlated metabolites identified from the PRISM IBD dataset are shown.

Additionally, within the IBD (PRISM) dataset, MiMeNet identified 351 well-predicted metabolites from the 466 annotated metabolites. Even though MelonnPan uses a default correlation cutoff of 0.3, when using the same correlation cutoff derived by MiMeNet, MelonnPan identified 198 well-predicted metabolites of which 181 were identified by MiMeNet. In the cystic fibrosis dataset, MiMeNet identified 143 well-predicted metabolites while MelonnPan identified 104. In the soil dataset, MiMeNet identified 29 well-predicted metabolites while MelonnPan identified 4. When training using the entire IBD (PRISM) dataset to predict the IBD (External) test data, MiMeNet identified 308 well-predicted metabolites while MelonnPan identified 186, of which 160 were also identified by MiMeNet (S2 Fig). When analyzing the overall prediction correlation and number of well-predicted metabolites, we observed that MiMeNet’s improvement was not a global improvement across all metabolites, but rather it came from MiMeNet being able to model a large set of microbes that MelonnPan could not. For example, in the IBD (PRISM) dataset, there were 237 metabolites with a negative prediction correlation. Of these metabolites, MiMeNet was able to predict 160 with a correlation above the determined cutoff. These metabolites also made up the set of metabolites with a prediction correlation of 0 in the IBD (External) set when using MelonnPan. Upon investigation, this set of metabolites was predominantly composed of triacylglycerols, long-chain fatty acids, and bile acids. These three classes of metabolites have been shown to interact with various microbes as well as relate to IBD patients. In addition, we observed that the running time of MiMeNet was robust and did not scale largely with the number of metabolites as all three datasets complete in similar timespans (Table 2). These results show that MiMeNet benefited from multi-task learning, the scalability of MLPNN, and the ability of MLPNN in capturing complex relationships between microbiome and metabolomes.

**Table 2.**
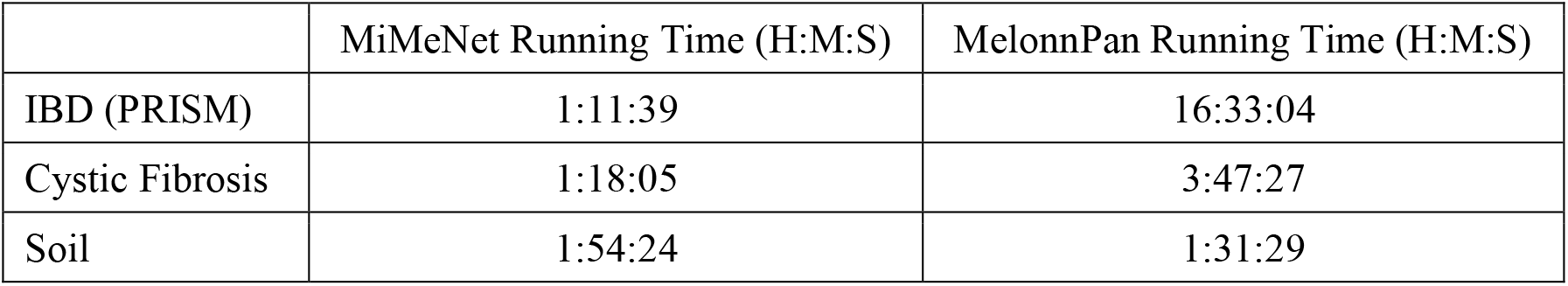
Running time for MiMeNet and MelonnPan. Running time for MiMeNet was calculated using the recommended settings from the Github page: 20 random searches for hyper-parameter tuning, 10-iterations of 10-fold cross-validation for prediction evaluation, 10-iterations of 10-fold cross-xvalidation for background generation, and the training of 10 final candidate models. MelonnPan was run using 10-iterations of 10-fold cross-validation and for the IBD (PRISM) datset MelonnPan only used the 466 annotated metabolomic features.

|  | MiMeNet Running Time (H:M:S) | MelonnPan Running Time (H:M:S) |
| --- | --- | --- |
| IBD (PRISM) | 1:11:39 | 16:33:04 |
| Cystic Fibrosis | 1:18:05 | 3:47:27 |
| Soil | 1:54:24 | 1:31:29 |

### MiMeNet identifies biologically important modules of microbes and metabolites

To discern what group of microbes contributed collectively to a group of metabolites, we computed the interaction scores for all pairs of microbes and the 6857 well-predicted metabolites using the network weights of the trained models obtained from the IBD (PRISM) data set. We identified 163 microbes that had at least one significant interaction with a well-predicted metabolite (Methods). A positive score means that the microbe contributes positively to the prediction of the abundance of the metabolite. Likewise, a negative score contributes negatively to the prediction of the abundance of the metabolite. To reveal the pattern of interaction scores, we grouped the microbes and metabolites into modules using biclustering (Methods). We identified 8 modules of microbes and metabolites respectively based on the biclustering analysis (Fig 5A) and computed the module feature value as the average abundance of features within the module for each sample. We then constructed a bipartite graph between microbe and metabolite modules such that the interaction score was the mean score found within the block identified by both modules (Fig 5B). To evaluate if the modules embody the useful information, we trained neural network models to predict patient IBD status based on either the module feature values or the original microbial and metabolite abuandance using the IBD (External) dataset. We observed that MiMeNet’s modules had similar predictive power to the original features (Fig 5C). Moreover, the metabolite abundance values and their module feature values are more predictive of IBD status compared to the microbial abundance and their module feature values, respectively. This result indicates that MiMeNet has extracted useful modules that are predictive to host disease status.

**Figure 5.**
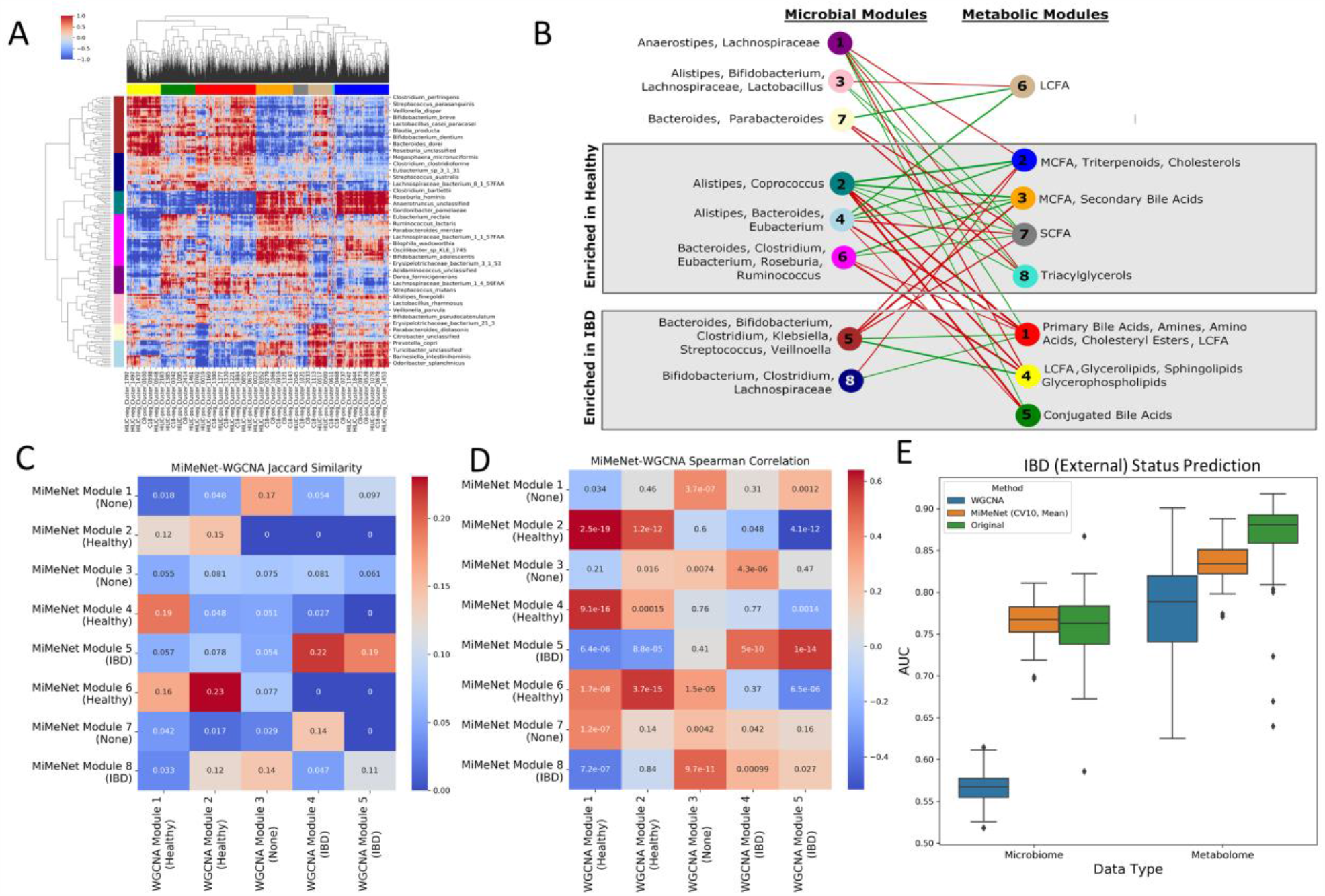
Clustering of microbes and metabolites in the IBD datasets. (A) Clustering of microbes (row) and metabolites (column) based on the feature importance scores given with Olden’s method. Row and column colors represent assigned modules. (B) Network connecting microbial modules with metabolomic modules. The most abundant genera are annotated for microbe modules. The most abundant metabolite classes are annotated for the metabolite modules. Red connections indicate negative interaction and green edges indicate positive interactions. Node color represents module color from (A). (C) Jaccard Index and (D) Spearman correlation between module features of WGCNA and MiMeNet. (E) IBD status prediction using WGCNA module feature values, MiMeNet module feature values, and original abundance from microbial and metabolomic data.

We further went to examine if a module is enriched for one patient group (IBD or healthy) by comparing the average normalized feature values of the members within the module between the two groups using the IBD (PRISM) samples (P<0.05, Wilcoxon rank-sum test) (S3 Fig). Four metabolite modules were enriched in healthy subjects, i.e., higher average module feature values. The first module (module 2) contained medium-chain fatty acids (MCFA), triterpenoids, and cholesterols. Triterpenoids, such as oleanolic acid and maslinic acid, have been shown to have an anti-inflammatory effect as well as enhance the integrity of intestinal tight junctions [36], and have been explored as therapeutic options for IBD [36-38]. In addition, both cholesterols and MCFAs have been noted to be depleted in subjects with IBD [39]. The second module (module 3) contained MCFA molecules as well as secondary bile acids such as deoxycholic acid and lithocholic acid, which was found to be reduced in IBD patients [40, 41]. The third module (module 7) was mainly composed of short-chain fatty acids (SFCA) such as propionate, butyrate, and valeric acid, all of which were shown to be protective against IBD [34]. The final module (module 8) contained triradylcglycerols, which in previous studies have been found depleted in subjects with IBD [39].

Similarly, three metabolite modules were found enriched in IBD patients. The first module (module 1) was composed of a large portion of primary bile acids, amines, amino acids, cholesteryl esters, and long-chain fatty acids (LCFA). Primary bile acids aid in the digestion of lipids and are further deconjugated to secondary bile acids by microbes in the gut. The deconjugation of primary bile acids in subjects with IBD was previously shown to have an impaired ability causing a greater abundance of primary bile acids[40]. Additionally, these primary bile acids were shown in previous studies to bind to the farnesoid X receptor, which is linked to the elevated immune response in IBD [42]. Cholesteryl esters have also been shown to be elevated in subjects with IBD, potentially explained by lipid mobilization or by increased intestinal permeability [43]. The amine group within this module is composed of N-acylethanolamines as well as sphingosine. N-acylethanolamines have been shown to alter the gut microbiome and potentially increase levels of lipopolysaccharides in the intestines, causing inflammation [44]. Sphingosine along with fatty acids that make up ceramide were also found within this module. Ceramide is a precursor to sphingosine-1-phosphate, a signaling sphingolipid that has been implicated in increased inflammation of the gut [45]. The LCFAs found here have been implicated with IBD contained eicosapentaenoic acid, arachidonic acid, and docosapentaenoic acid, which were all identified in the original study as well [15]. The largest group in this module, however, comprised of 31 (24.4%) different amino acids and amino acid derivatives. Amino acids have been found in a previous study to be elevated in IBD patients, possibly due to the increase in the bacterial enzyme urease promoting a flux of nitrogen for the synthesis of amino acids by the host [46]. The second module (module 4) contained LCFAs such as docosahexaenoic acid, arachidonic acid, and eicosapentaenoic acid, as well as glycerolipids, glycerophospholipids, and sphingolipids. Bacterial-derived sphinogolipids have been shown to play a crucial role in the development of IBD through multiple signaling pathways [47]. The elevated glycerolipids and glycerophospholipids were also identified in the original study [15]. The third module (module 5) contained mostly conjugated bile acids, which is similar to primary bile acids have been shown to be elevated in IBD subjects due to a decreased ability to deconjugate the bile acids into secondary bile acids. In addition to disease related metabolite modules, MiMeNet also identified one module (module 6) not associated with disease status. This module was composed of a small number of LCFAs.

Lastly, we compared the microbial modules in the IBD (PRISM) dataset identified by MiMeNet to those identified by the Weighted Correlation Network Analysis (WGCNA) (Methods). The module features of a sample were calculated as the average normalized abundance of the members within the module. Using the Jaccard similarity between the members of the modules as well as the Spearman correlation coefficient between module features across samples, we observed only a small consensus between the two groupings (Fig 5 (C,D)). Furthermore, the evalution of prediction of IBD status on the IBD (External) dataset by the neural network models trained using the feature values of the WGCNA modules had a much lower mean AUC value (0.558) compared to those generated from the original microbial abundance and the feature values of the MiMeNet modules (Fig 5E). Additionally, we compared the prediction of IBD status when MiMeNet uses k-fold cross-validation (k=10, 5, 3, and 2) and observed that modules generated using 10-fold cross-validation were most predictive of IBD status (S4 Fig). Taken together, the analyses demonstrate that MiMeNet has organized microbes and metabolites into biologically meaningful modules.

### MiMeNet identifies biologically important microbe-metabolite module interactions

We analyzed how the microbe modules impacted the various metabolite modules. SCFAs, and in particular butyrate, have been shown to be protective of inflammation in the gut and important for gut homeostasis [48]. A recent study identified ten microbial species commonly found in the gut microbiome that produce butyrate [49]. Five of these microbes, *Eubacterium biforme, Eubacterium hallii, Eubacterium rectale, Roseburia intestinalis*, and *Roseburia inulinivorans* were found in microbial module 6, which had a strong positive association with the metabolite module comprising of SCFAs. For comparison, we calculated the Spearman correlation p-value between butyrate and each microbe in the dataset and selected significant pairs after multiple test correction (adj.P<0.05, Benjamini-Hochberg). Although the pairwise analysis identified 12 significant correlations, only one pair contained a microbe, *Anaerostipes hadrus*, from the set of butyrate producing microbes, indicating the pairwise univariate analysis can only capture the strongest correlation pairs. This suggests that not only could MiMeNet capture more biologically relevant microbe-metabolite interactions but that it can also group them into meaningful modules based on common interactions.

In addition, we observed that metabolite module 5 had very weak positive interactions and very strong negative interactions with microbial modules 2 and 4. This module contained primarily conjugated bile acids, which are formed when primary bile acids are conjugated with either taurine or glycine in the liver. Since this process is performed independently of the gut microbiome, it would be expected to not observe a strong positive association. However, once these bile acids enter the gut, various microbial enzymes convert these conjugated bile acids into secondary bile acids through deconjugation, oxidation, and 7-dehydroxylation. Previous studies have identified multiple genera of microbes that produce enzymes responsible for this conversion, including *Bacteroides, Clostridium, Eubacterium*, and *Ruminococcus* [50], all of which constitute a large portion of microbial modules 2, 4, and 6. These modules all show a negative interaction with conjugated bile acids and a positive interaction with secondary bile acids, suggesting that microbes found within these modules regulate the conversion of conjugated bile acids to secondary bile acids.

## Discussion

We have developed MiMeNet for integrative analysis of the paired microbiome-metabolome datasets using neural networks. The objectives of MiMeNet are to model microbiome-metabolome interactions and uncover functional relationships between microbes and metabolites. Using datasets obtained from the human gut of subjects with IBD, lung sputum of subjects with cystic fibrosis, and soil wetting environmental studies, we showed that MiMeNet models can produce a robust and accurate prediction of the community metabolome based on the microbial taxa abundance in both cross-validation and external validation. MiMeNet is empowered by neural networks’ capability of modeling nonlinear relationships and multi-task learning, which predicts the abundance of all metabolites, a set of homogenous tasks. The entire set of metabolite outputs can be considered sets of sub-components from multiple metabolic and physiological pathways, which have varying functional impacts on the host or environment. We consider these sets of metabolites as different tasks, and due to the hierarchical nature of pathways, there are often shared metabolites between pathways that may facilitate prediction through shared information, although the loss of each task was weighted equally in the loss function. Our results of the IBD data demonstrated that the MiMeNet prediction on the set of the annotated metabolites benefited from including tasks of predicting the rest of the unannotated metabolites in the metabolome profiles (Fig 2(D-F)). We also compared the performance of the prediction using two types of microbial abundance representations: relative abundance (RA) and the centered log-transformation of abundance (CLR). The prediction correlations in the IBD (PRISM) dataset were comparable between both transformations, however, we saw an increase in correlations in the cystic fibrosis and soil datasets when using CLR. In addition, we observed an improvement in prediction performance on the IBD (External) test set when using the CLR transformation (S5 Fig). Moreover, there was a large consensus of well-predicted metabolites using both transformations, but MiMeNet was able to identify slightly more well-predicted metabolites when using CLR (S6 Fig). We also observed that the modules generated using CLR were more predictive of IBD status than those generated using RA values (S4 Fig). We note that since not all metabolites may be associated with microbes, they will present a lower prediction correlation that results in an overall lower mean correlation across all metabolites. MiMeNet filters out these metabolites by using a threshold generated from an empirical background distribution of correlation values from the bootstrapping procedure. This limitation may be alleviated by integrating other omics data such as metatranscriptomic data in the future. We also observed a higher threshold value for the soil data. This may be due to the small sample size and as well as the longitudinal observations, which may require an elaborated data shuffling scheme in the bootstrapping procedure.

MiMeNet also facilitates additional analysis from the learned network models. Using the IBD data, we showed that the interaction scores derived from the network weights could be used to construct modules of microbes with similar positive or negative effects on a set of metabolites. Grouping metabolites into modules with similar interaction patterns and functions can facilitate the annotation of uncharacterized metabolites through “Guilt of Association” [15, 51]. This is extremely important in the field of metabolomics due to a large amount of current “dark matter” [52]. Untargeted metabolomics study has been hindered by many unknown metabolites and much of the newly collected information remains uninterpreted. The annotated metabolites clustered in a MiMeNet module that contains unannotated metabolites may provide a clue that those metabolites participate in similar biochemical reactions due to similar interaction patterns with some microbial modules. Those unannotated metabolites may relate to the annotated metabolite structurally or functionally. Additionally, if microbial gene features are used as input to train MiMeNet networks, it may further uncover gene-metabolite association from the interaction scores identified from the MiMeNet model. Although the MiMeNet analysis is data-driven without incorporating mechanistic knowledge. Nevertheless, these types of evidence obtained from the integrative analysis of metagenomes and metabolomes could be used in predictive computational approaches such as MAGI and MINEs to increase the confidence of metabolite identification [53]. Furthermore, we have shown that the predictive models learned from the modules were equally competitive for the prediction of the host IBD status compared to those built on the entire microbes or metabolites, suggesting the metabolic functional relevance of microbial modules. This direction could be further explored in integrating omics data for host phenotype prediction [23].

The predictive model in MiMeNet distinguishes from MelonnPan [26], which uses a regularized linear regression to model each metabolite separately. MiMeNet models all metabolites nonlinearly and benefits from learning of the shared information. The MiMeNet model is also different from another highly relevant predictive model of NED in several aspects. NED models the metabolome using the representation of microbiome in the latent space generated by an encoder [28, 29]. It imposes an additional sparsity constraint to prevent overfitting and non-negative weights for ease of interpretation. Our comparison shows that the MiMeNet model is competitive with the NED model in terms of predictive accuracy and the ability to make well-prediction for a larger set of metabolites (S7 Fig). This suggests that the constraint of non-negative weights may diminish the capability of the NED to capture interactions where some microbes negatively affect some metabolites. Indeed, it was reported that more than 50% of associations identified between microbial metabolic pathways and species in human fecal and blood samples are negative [19]. Similar to MiMeNet, the NED generated latent microbiome space, which was shown to contain biologically relevant information useful in discrimination of Crohn’s disease, ulcerative colitis, and healthy subject, and prediction of other clinical measurements such as usage of immunosuppressive therapy. However, since NED models metabolites using the microbiome latent space representation, the interpretation of microbe-metabolite interaction is less intuitive, and it does not reveal modules in microbes and metabolites with shared interaction patterns. Nevertheless, we note that NED belongs to a broad class of integrative approaches to omics data analysis using latent components derived from various statistical and machine learning models [54].

The scope for integrative analysis of microbiome and metabolomics data developed so far are diverse, ranging from the identification of statistical associations using univariate and multivariate correlation analyses to predictive modeling based on machine learning and metabolic network-based modeling [15, 18, 55-59]. For example, a recently released webserver, M2IA [60], is an excellent tool to streamline the statistical analysis, such as overall similarity assessment of the two omics datasets, pairwise correlation analysis between microbes and metabolites, a heatmap visualization tool for revealing positive and negative correlation patterns. It also includes other components including WGCNA for identifying clusters of metabolites based on the pairwise Spearman’s correlations of metabolites, supervised multivariate analyses of detection of disease associations on the microbes, or metabolites or functional annotations. M2IA, however, does not provide predictive models to quantify the impact of the abundance of multiple microbes on a metabolite abundance. This is the major distinction between MiMeNet and other machine learning-based methods (MelonnPan [26], mmvec [27], and NED [28, 29]) mentioned early. Despite the progress, all these methods including MiMeNet, are still limited in providing biological plausibility and mechanistic insights. Future direction for MiMeNet extension could be to develop procedures to discern the statistical association of microbe-metabolites interactions to host diseases and detect modules of metabolites that require specific microbial taxa in healthy and disease samples. In addition, datasets including additional metagenomics data could be integrated to allow for predicting metabolites from genes. This could potentially reveal richer functional connections that are not currently annotated as well as provide a richer functional exploration of the impact of the microbiome on the host metabolome. Moreover, neural networks could be utilized to model how the host metabolome influences gut microbial composition.

## Methods

### Data for Evaluation

Concurrent profiles of microbiome and metabolome from varying environments were used for evaluation (Table 1). The first dataset was taken from a published study of patients with inflammatory bowel disease (IBD) [15]. It includes one cohort from the Prospective Registry in IBD Study at MGH (PRISM), which enrolled patients with a diagnosis of IBD based on endoscopic, radiographic, and histological evidence of either Crohn’s Disease (CD) or Ulcerative Colitis (UC). This dataset has 121 IBD patients and 34 controls, and is named as IBD (PRISM). Additionally, it includes an external validation dataset using two other cohorts. One consists of 20 healthy subjects who participated in LifeLines-DEEP, a general population-based study in the northern Netherlands (NLIBD) [61]. The second cohort consists of 43 subjects with IBD taken from the Department of Gastroenterology and Hepatology at the University Medical Center in Groningen, Netherlands. This dataset is named IBD (External). The processing of the stool samples collected is described in the original study [15]. A total of 201 microbial species and 8848 metabolites were identified for the IBD (PRISM) and IBD (External) datasets.

The second dataset was taken from a study that collected 172 lung sputum samples from patients with cystic fibrosis [31]. Microbial features were generated using 16S rRNA gene sequencing and abundance was collected at the genus level, resulting in 657 unique microbial features. Metabolomic data were generated using LC-MS/MS technology, resulting in 168 unique metabolites.

The third dataset represents microbial and metabolic activity caused by soil wetting at five-time points across four biocrust successional stages [32]. Biocrust soil water for each sample was analyzed by LC/MS for metabolite detection. Metagenomic shotgun sequencing was used to profile the microbial community and the authors used the 50S ribosomal protein L15 to map microbial taxa. A total of 466 microbes and 85 metabolites were detected.

Any input or output feature that is present in less than 10% of samples was removed. Microbiome and metabolomic data were then transformed using the Center log-ratio (CLR) transformation:

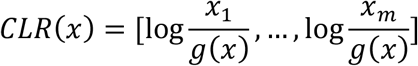

where ***x*** is the abundance vector of a sample and *m* is the number of features. The only exception was for the IBD (PRISM) and IBD (External) microbe values, which were obtained in relative abundance (RA).

### MiMeNet Architecture and training protocol

An MLPNN model is composed of multiple fully connected hidden layers composed of perceptrons. The values *h*_*l*+1_of layer *l* + 1are defined as

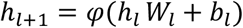

Here *W*_*l*_ are the weights connecting the perceptrons of the *l*^*th*^ layer with values *h*_*l*_ with those of layer *l* + 1, *b*_*l*_ is the bias values between layer *l* and *l* + 1, and *φ* is a non-linear function. In MiMeNet, *φ* is set as the rectified linear unit (ReLU). We selected this activation function since previous studies have shown that it is resilient to the problems of exploding and vanishing gradient in MLPNN training [62]. We used the L2 regularization in order to prevent the model from having large weights. Additional regularization was applied through dropout at each hidden layer, where a portion of the nodes and their weights are masked for a given epoch. MiMeNet was trained using the ADAM optimizer and the mean squared error (MSE) loss function. The entire loss function for MiMeNet is defined as,

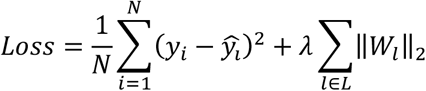

Here, *N* is the number of training samples and *L* is the total number of hidden layers. The first term represents the mean squared error (MSE) of the observed metabolites *y* and the predicted metabolites *ŷ*. The second term represents the L2-regularization penalty which is controlled by the parameter *λ*.

The overall evaluation of MiMeNet prediction was conducted using 10 iterations of the 10-fold cross-validation, and the average of the correlations between the predicted and observed values for metabolites was reported. More explicitly, during the 10-fold cross-validation, each dataset was partitioned into two subsets: 90% for training and 10% for testing. For each training partition, the 90% of the data was further split into 80% for model training and 20% for validation. After finishing one iteration of 10-fold cross-validation, the Spearman’s correlation coefficient (SCC) between the predicted and the observed was calculated for each metabolite. To prevent overfitting, MiMeNet models were trained using early stopping. After each iteration of updating network weights using the 80% of the training set, the loss of the validation set was calculated. The training process was terminated when the loss of the validation set has not improved within 40 iterations, and the network weight parameters were set to the values of the best performing model on the validation set. Finally, the average of the SCC values was calculated after repeating the 10-fold cross-validation procedure for 10 times. For the IBD datasets, a final model trained on the full IBD (PRISM) dataset was then evaluated on the IBD (External) test set.

Hyper-parameter tuning was performed on the first training partition during cross-validation. To determine the optimal set of hyper-parameters (number of layers, layer size, *λ*, and dropout rate), we performed a cross-validated random search using a nested 5-fold cross-validation. We allowed for 1, 2, and 3 hidden layers of sizes 32, 128, and 512. The L2 regularization parameter (*λ*) was selected from 10 different values between 0.0001 and 0.1, evenly spaced on a log scale. Dropout was selected from 0.1, 0.3, and 0.5. The average Spearman correlation coefficient (SCC) was calculated after a model was trained. We evaluated 20 sets of hyper-parameters and selected the best performing set for the rest of the 10-fold cross-validataion. The optimal hyper-parameters are shown in Table 3.

**Table 3.**
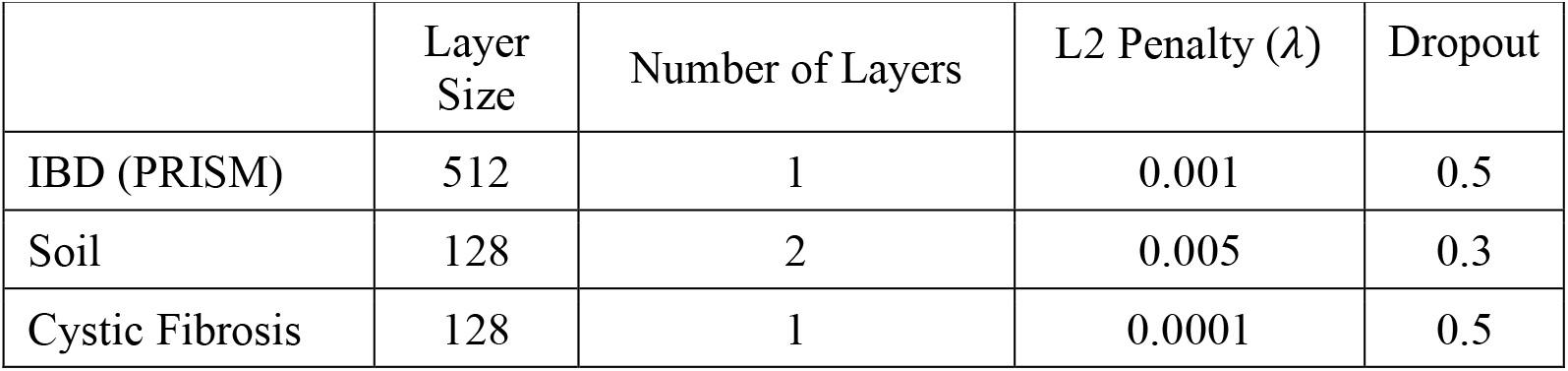
The optimal hyper-parameters for the neural network model in each dataset.

### Identifying Well-predicted Metabolites

To identify which metabolites are well predicted by MiMeNet, we generated a background distribution of SCC by training 100 models of 10-fold cross-validation where the samples in microbiomes and metabolomes were each randomly shuffled. From each of the 100 models generated, we collected the prediction SCC for each metabolite and combined the values across all metabolites to construct the background distribution. We then defined a metabolite as well-predicted if its SCC is above the 95th percentile of the background correlations.

### Calculation of Microbe-Metabolite Interaction Scores

Microbe-metabolite interaction scores are calculated using Olden’s method for understanding variable contributions in neural network models [63]. Olden’s method generates these scores by multiplying the weights of each hidden layer together, resulting in a single matrix where each row represents an input feature and each column represents an output feature. More specifically, a microbe-metabolite interaction matrix S is calculated as

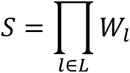

where *l* is the current layer in the set of *L* layers, and *W*_*l*_ is the weight matrix connecting layer *l* − 1and layer *l*. Each element in *S* represents a microbe-metabolite interaction score. A positive value indicates that an increase of the microbe will lead to an increase of the metabolite and a negative value indicates that an increase of the microbe will lead to a decrease of the metabolite. To identify microbes with significant interactions, we extracted interaction matrices from the network models used to generate the background correlation distributions and calculated the mean interaction matrix. We then filtered only the well-predicted metabolites in both the observed and background interaction matrices. The background matrix was then flattened into a vector and a threshold was set at the 97.5 percentile. Any observed interaction score with an absolute value above the threshold was considered significant. Lastly, any microbe without significant interaction with a metabolite was filtered from the interaction score matrix.

### Clustering and Visualization of Microbe-Metabolite Interactions

Before biclustering we normalized interaction scores by dividing each value by the significant threshold score identified by the background and clipped values to be between 1 and -1. In doing so, every significant interaction was treated with equal magnitude and non-significant interactions were scaled between 1 and -1. Microbe-metabolite interaction scores were used to cluster microbes and metabolites separately based on the Euclidean distance and complete linkage using Seaborn’s *clustermap* function in Python. Modules were constructed by cutting each dendrogram at a given height. To determine the number of clusters for microbes and metabolites, we generated a consensus matrix using the trained cross-validated models [64]. For each model, the interaction score matrix was filtered to only include well-predicted metabolites and microbes with significant interactions. The resulting matrices were clustered on row and column separately where the number of clusters, *k*, ranged from 2 to 20. The consensus matrix for *k* clusters, *M*^(*k*)^, was calculated as the mean connectivity matrix across all clusterings,

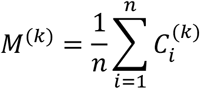

where 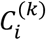 is the connectivity matrix of the clustering using *k* clusters on the *i*^*th*^ model, and *n* is the total number of trained models. We then calculated the area under the cumulative distribution function (CDF) for the consensus matrix of each clustering,

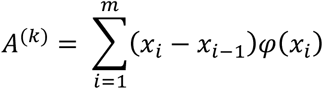

where *x* is a set of locations along the x-axis and *φ*(*x*_*i*_) is the CDF value at the position *x*_*i*_. For our analysis we used *x* = {0.01, 0.02, 0.03, …, 0.99,1.0}. Then we calculated the proportional change in area as the number of clusters changed,

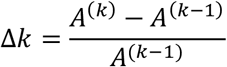

This value represents how much cleaner the consensus matrix gets if we increase the number of clusters by 1. We set a threshold of Δ*k* = 0.025 for both microbe and metabolite clustering, indicating that increasing the cluster by 1 more would give less than a 2.5% increase in the area under the CDF. An example is shown in S8 Fig.

For visualization of microbe-metabolite interaction networks, the average normalized interaction score between two modules was used as the mean score between each pair of microbes and metabolites within each module. For visualization purposes only we removed any interaction score whose absolute value was less than 0.25. Networks showing microbial and metabolite modules and the interactions between them were constructed using Cytoscape [65].

### Determination of the final model for external metabolite prediction

For evaluating the external test data, an ensemble of models is trained on the entire training dataset. For each trained model, clustering is performed, and microbe and metabolite consensus scores are assigned as the number of co-clustered microbes or metabolites that also occur in the clustering determined through the cross-validation consensus clustering. The model with the highest combined consensus scores was selected as the final model for external evaluation.

### Weighted Correlation Network Analysis (WGCNA)

WGCNA analysis of microbial features was performed using the *WGCNA* library in R [66] on each dataset. The adjacency matrix was created with a power of 2 for the microbiome data and 5 for the metabolomic data. To compare the WGCNA microbial modules to the MiMeNet microbial modules derived from the IBD (PRISM) dataset, we calculated the Jaccard similarity between the modules as well as the Spearman correlation.

### Neural networks for host phenotype prediction

For each module derived from MiMeNet or WGCNA using the IBD (PRISM) dataset, the module feature value was calculated as the mean normalized abundance value of the features comprising the module for each subject in the IBD (External) dataset. Network models for the IBD status prediction were trained using the original microbial OTUs (using either relative or center log-transformed abundance), the module feature values of the MiMeNet and WGCNA microbial modules, respectively. Similarly, the network models were trained using the normalized metabolomic abundance and metabolomic module feature values. We trained a 3-layer neural network model with 32 nodes in each layer for each input format described above. The evaluation was done by training 100 neural network models on the IBD (PRISM) dataset and evaluating on the IBD (External) test set using the area under the receiver operating characteristic curve (AUC).

## Other Existing Tools

The MelonnPan was downloaded from https://github.com/biobakery/melonnpan and executed using the given instructions. The NED model was trained using code downloaded from https://github.com/vuongle2/BiomeNED.

## Data Availability

The IBD amplicon and metabolomic abundance data for the IBD cohorts were obtained from the supplemental section of the original study [15]. BIOM files for the cystic fibrosis and soil datasets were obtained from the GitHub repository for mmvec (https://github.com/biocore/mmvec). All data is provided at https://github.com/YDaiLab/MiMeNet as well.

## Code Availability

Data and source code are available at https://github.com/YDaiLab/MiMeNet.

## Acknowledgements

We gratefully acknowledge the support of NVIDIA Corporation with the donation of the Titan Xp GPU used for this research.

## Author Contributions

Author Contributions Conceptualization: Derek Reiman, Yang Dai.

Data curation: Derek Reiman.

Formal analysis: Derek Reiman, Yang Dai.

Funding acquisition: Yang Dai.

Investigation: Derek Reiman, Brian T. Layden, Yang Dai.

Methodology: Derek Reiman, Yang Dai.

Project administration: Yang Dai. Resources: Yang Dai.

Software: Derek Reiman.

Supervision: Yang Dai.

Validation: Derek Reiman, Yang Dai.

Visualization: Derek Reiman.

Writing – original draft: Derek Reiman, Yang Dai.

Writing – review & editing: Derek Reiman, Brian T. Layden, Yang Dai.

## Competing Interests

The authors declare no competing interests.

## Materials & Correspondence

Requests for materials and correspondence should be addressed to.

## MiMeNet: Exploring Microbiome-Metabolome Relationships using Neural Networks

Derek Reiman, Brian T. Layden and Yang Dai

## Supplemental Figures

**S1 Fig .**
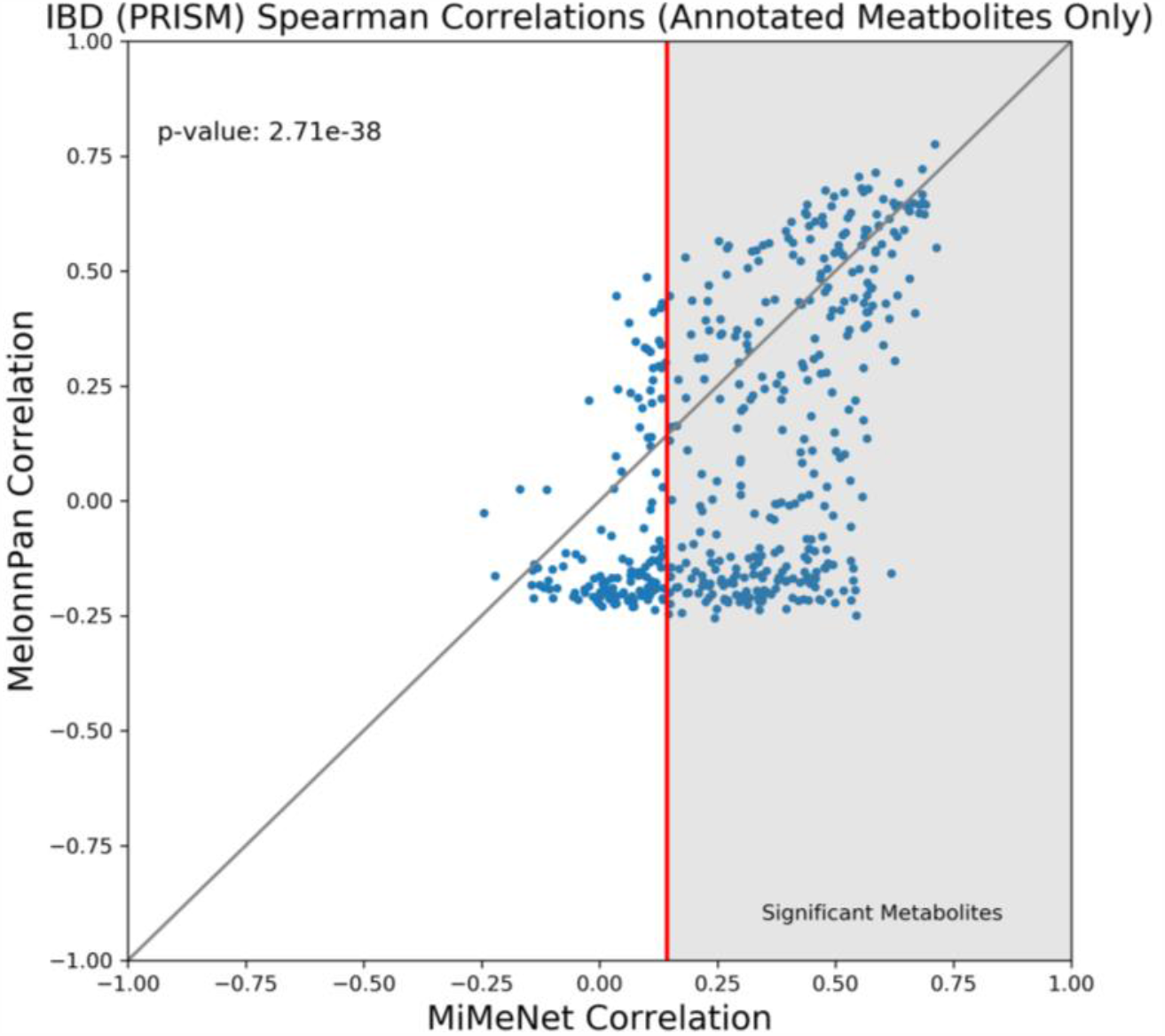
Prediction correlation comparison between models of MiMeNet and MelonnPan trained only on the annotated metabolites. Scatterplot of mean predicted Spearman’s correlations over 10 iterations of the 10-fold cross-validation for each metabolite between MiMeNet and MelonnPan when MiMeNet is only trained on the annotated metabolites in the IBD (PRISM) dataset.

**S2 Fig .**
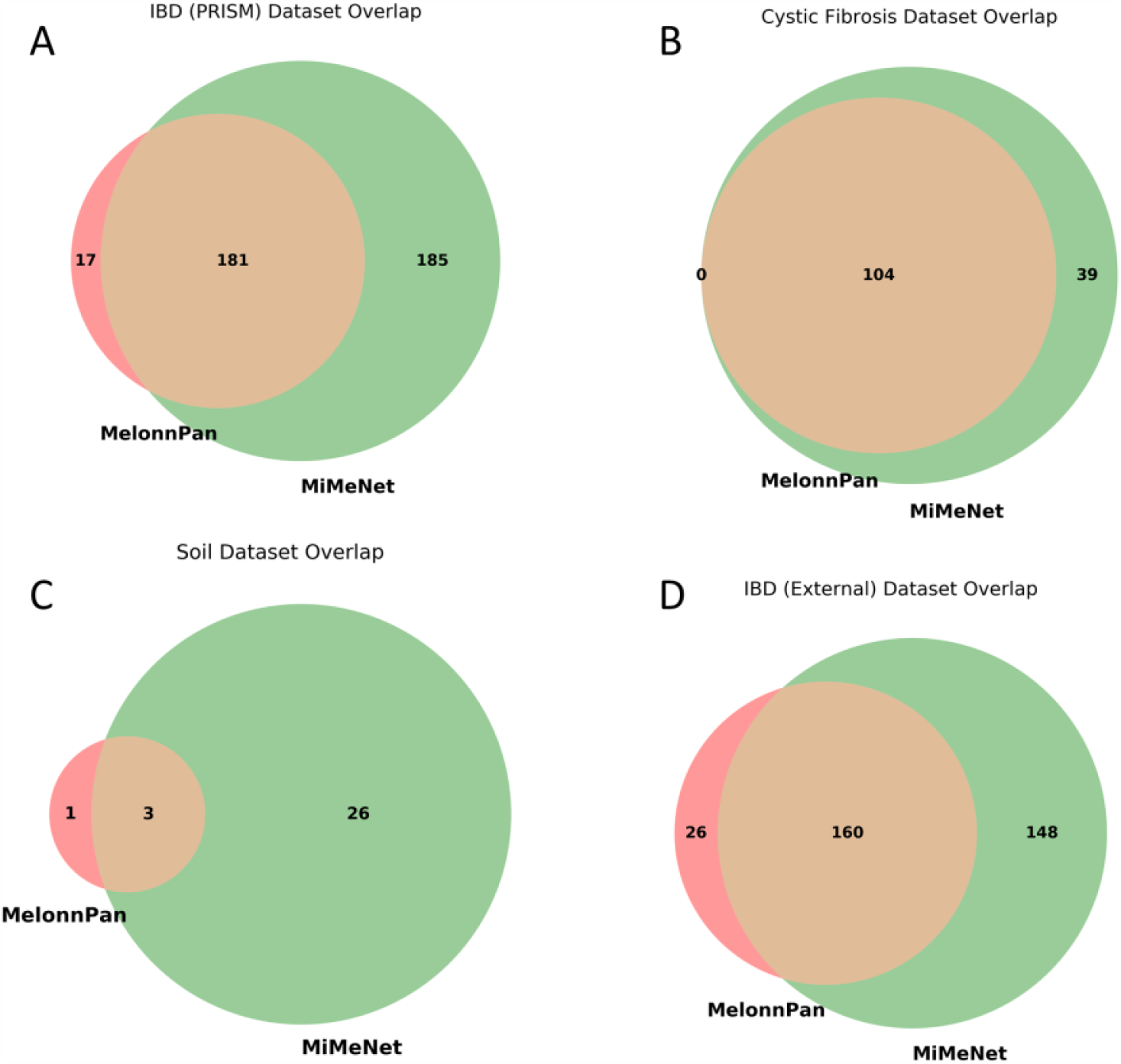
Overlap of well-predicted metabolites identified by MiMeNet and MelonnPan. Using the correlation cutoff identified by MiMeNet, the overlap between the well-predicted metabolites is shown between MiMeNet and MelonnPan for (A) IBD (PRISM) dataset, (B) cystic fibrosis dataset, (C) and soil dataset. (D) In addition, the overlap of well-predicted metabolites is shown when training on the entire IBD (IBD) dataset and predicting on the IBD (External) validation dataset.

**S3 Fig .**
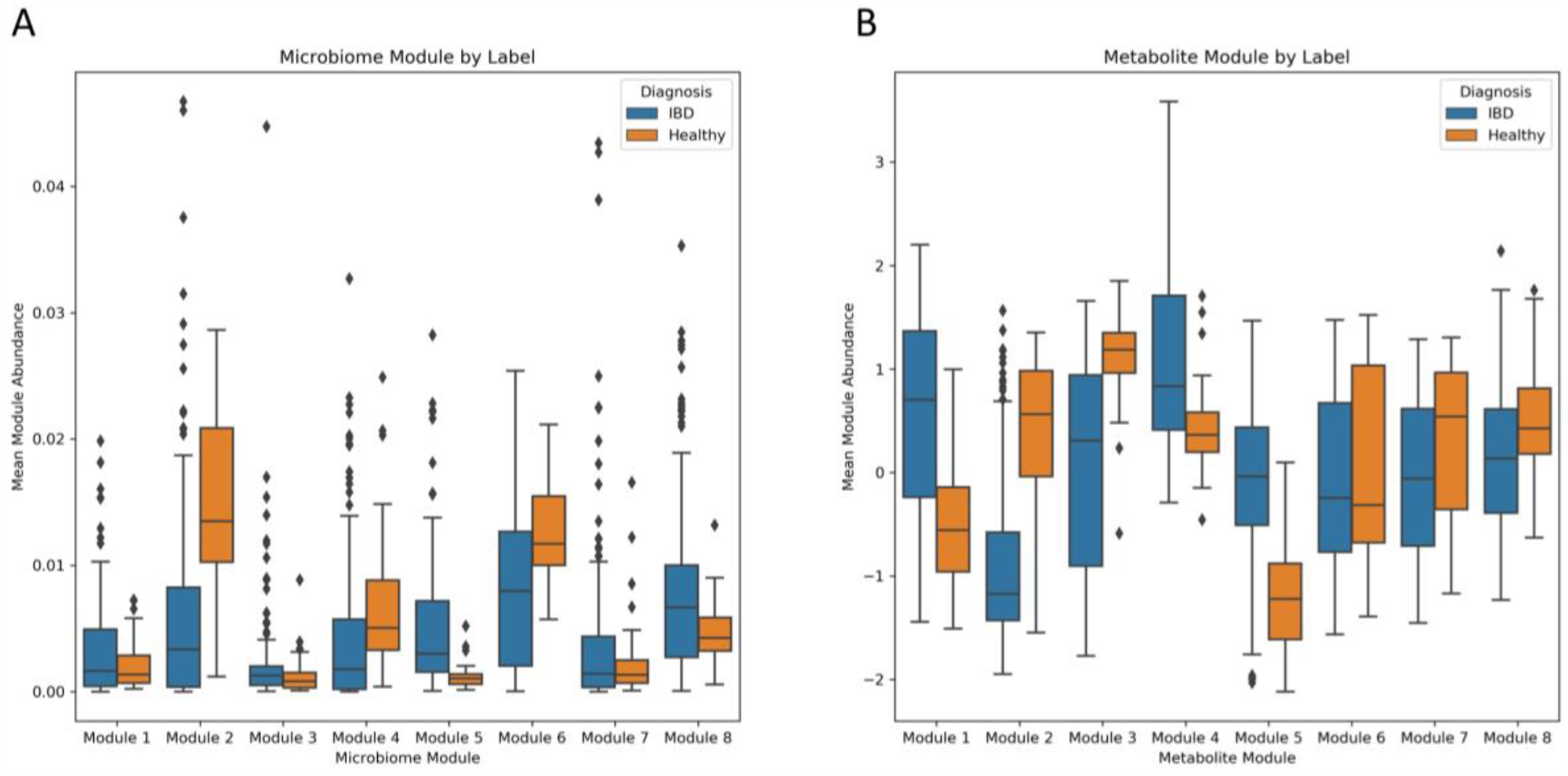
Microbial and metabolic module abundance by patient status in the IBD (PRISM) dataset. The mean abundance of members within a module are shown here for healthy patients and IBD patients using (A) microbial and (B) metabolic modules identified by MiMeNet.

**S4 Fig .**
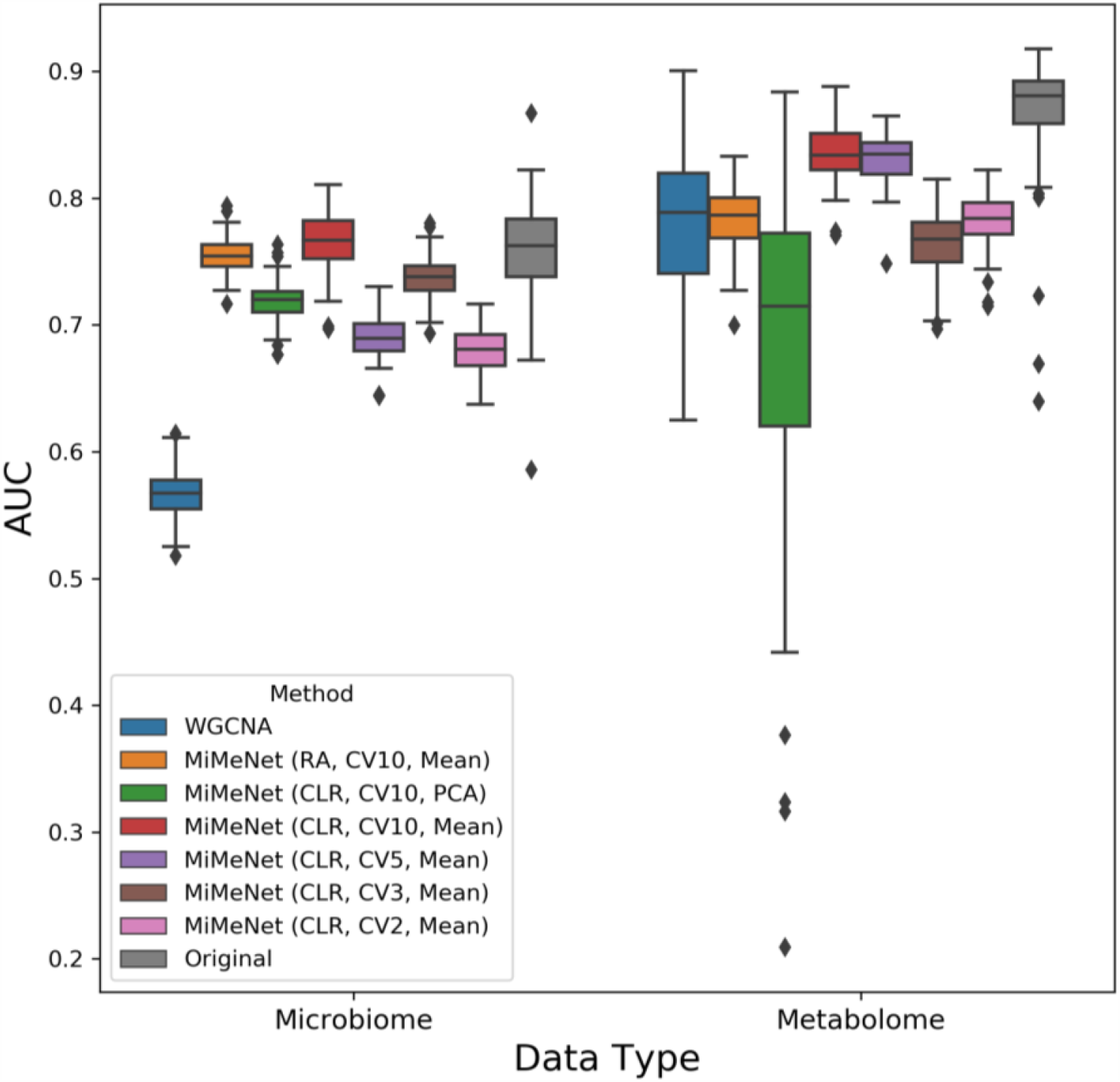
IBD status of prediction for different module construction methods evaluated on the IBD (External) dataset. Module values were constructed using WGCNA and MiMeNet. For metabolomic modules constructed by MiMeNet, the values within the parentheses represent the compositional transformation, number of folds for cross-validation, and the aggregation method respectively. Mean aggregation calculates the mean normalized abundance value. PCA aggregation uses the first principal component of the members’ values from that module. Microbiome modules were constructed similarly with the exception that RA was always used.

**S5 Fig .**
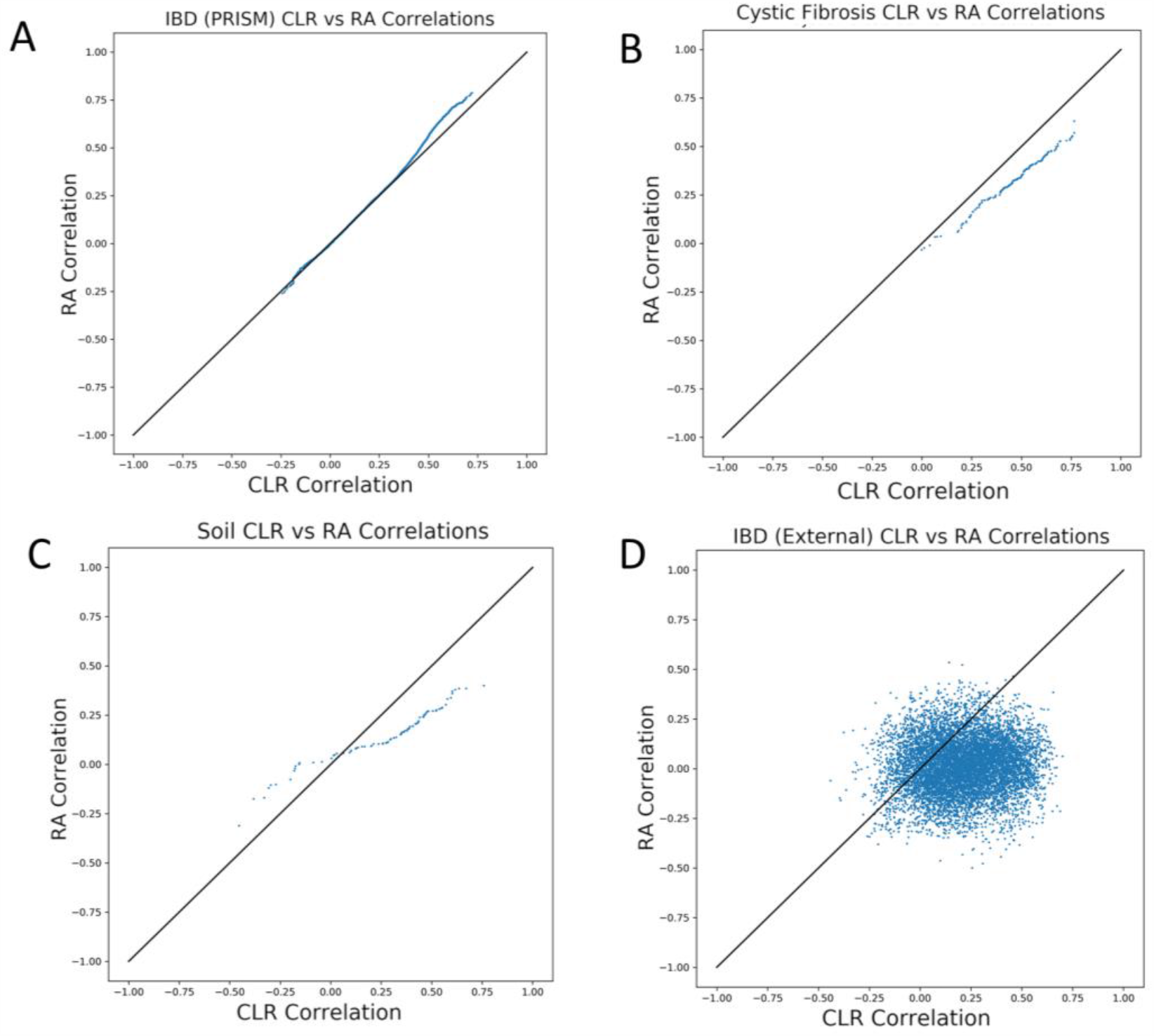
Comparison of prediction correlation when using relative abundance and centered log-ratio. Scatterplots comparing metabolite correlation prediction between data transformed to relative abundance (RA) and centered log-ratio (CLR) for (A) IBD (PRISM), (B) cystic fibrosis, (C) soil datasets using 10 iterations of 10-fold cross-validation, and (D) IBD (External) test predictions using models trained on the full IBD (PRISM) dataset.

**S6 Fig .**
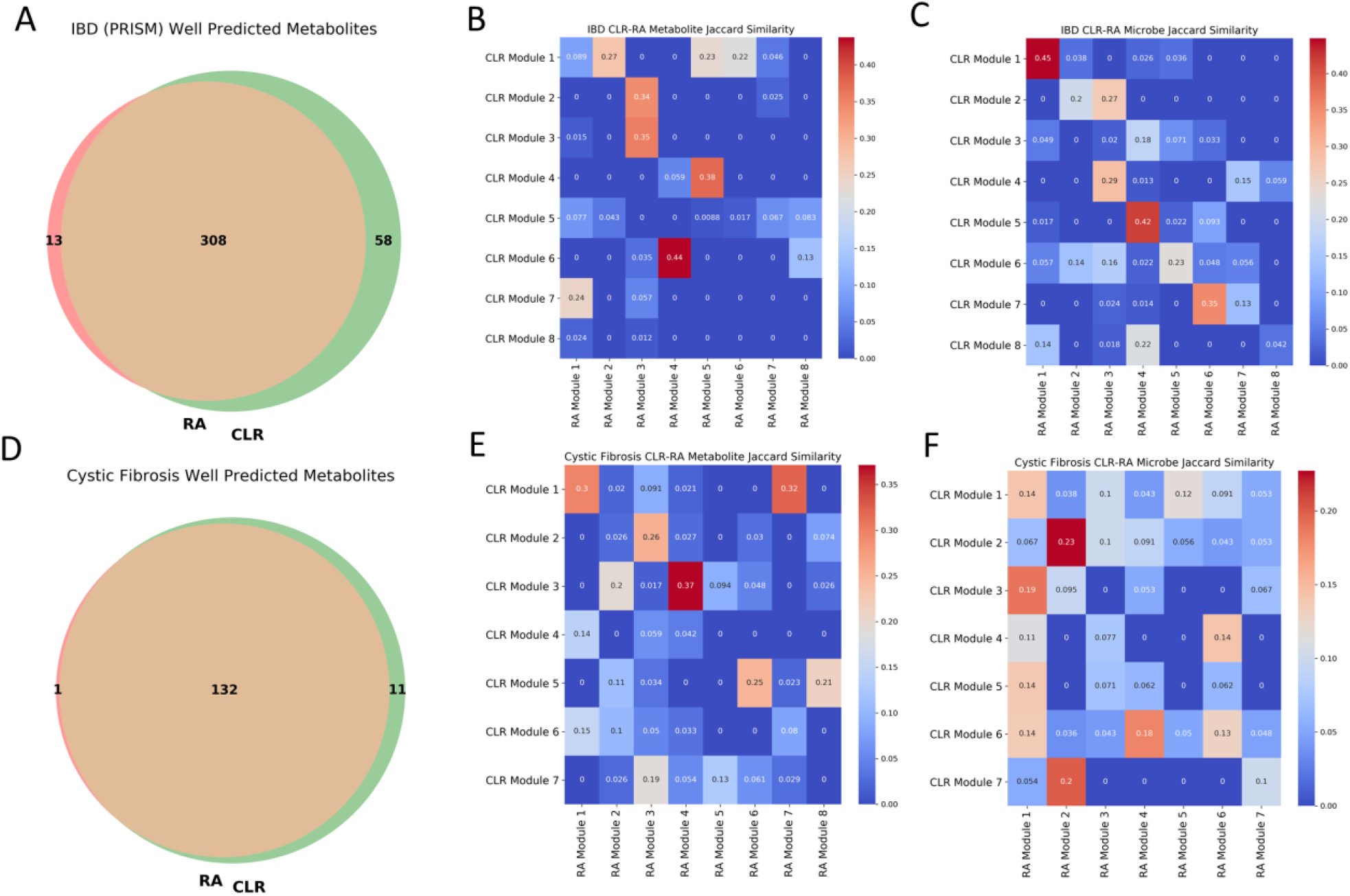
Overlap of significant metabolites and module membership when using relative abundance and centered log-ratio. (A) Overlap of well-predicted metabolites when using relative abundance and centered log-ratio for the IBD (PRISM) dataset. Heatmaps show Jaccard similarity between membership of (B) metabolite and (C) microbial modules in the IBD (PRISM) dataset. (D) Overlap of well-predicted metabolites when using relative abundance and centered log-ratio for the cystic fibrosis dataset. Heatmaps show Jaccard similarity between membership of (E) metabolite and (F) microbial modules in the cystic fibrosis dataset.

**S7 Fig .**
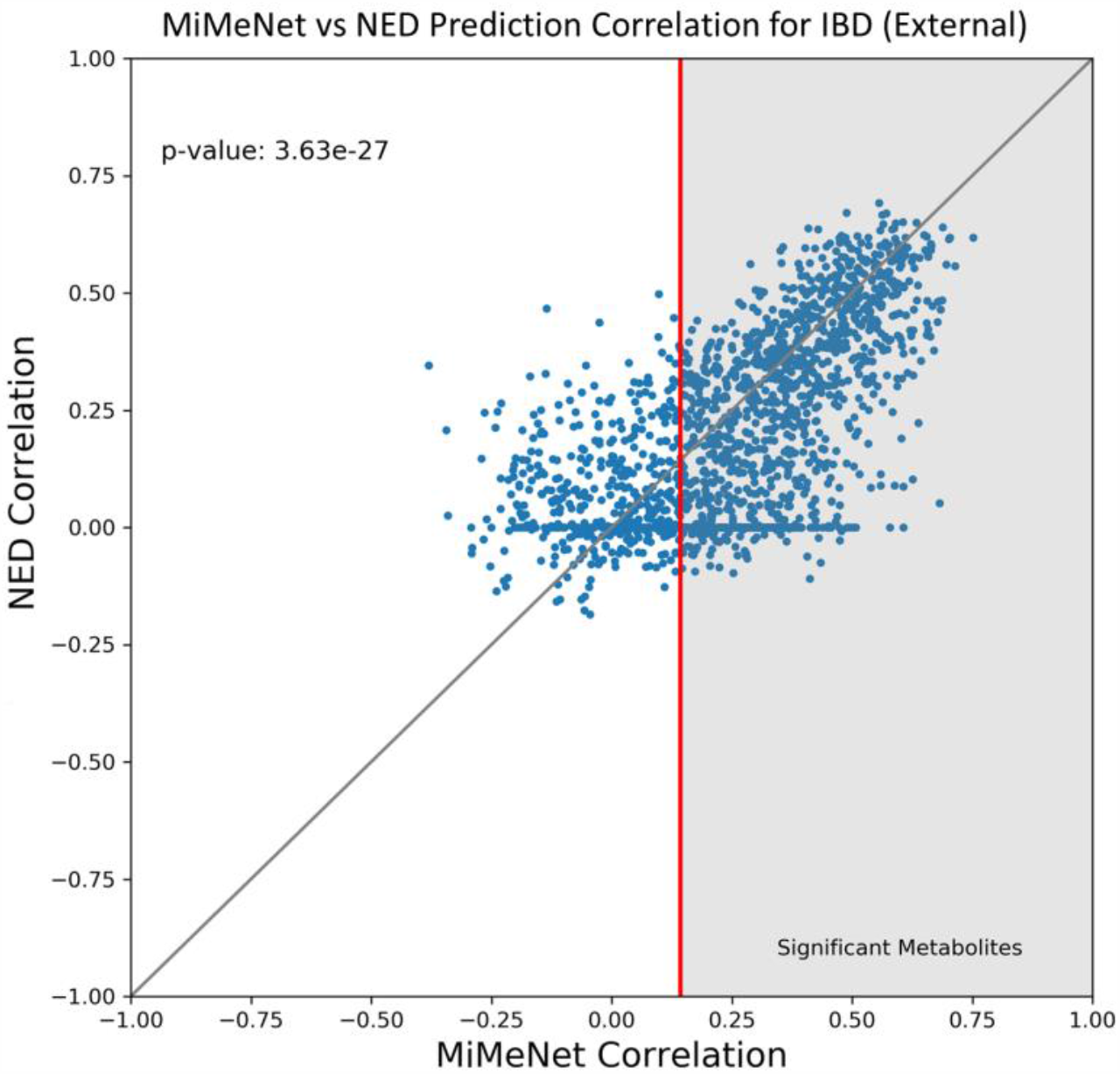
Comparison of MiMeNet to NED. Scatterplot comparing metabolite prediction correlation between MiMeNet and NED on the IBD (External) dataset validation. The red line indicates the correlation threshold identified by MiMeNet and the gray area represents well-predicted metabolites. The one-tailed p-value (Wilcoxon sign-rank) comparing MiMeNet’s values to NED’s is shown in the top right.

**S8 Fig .**
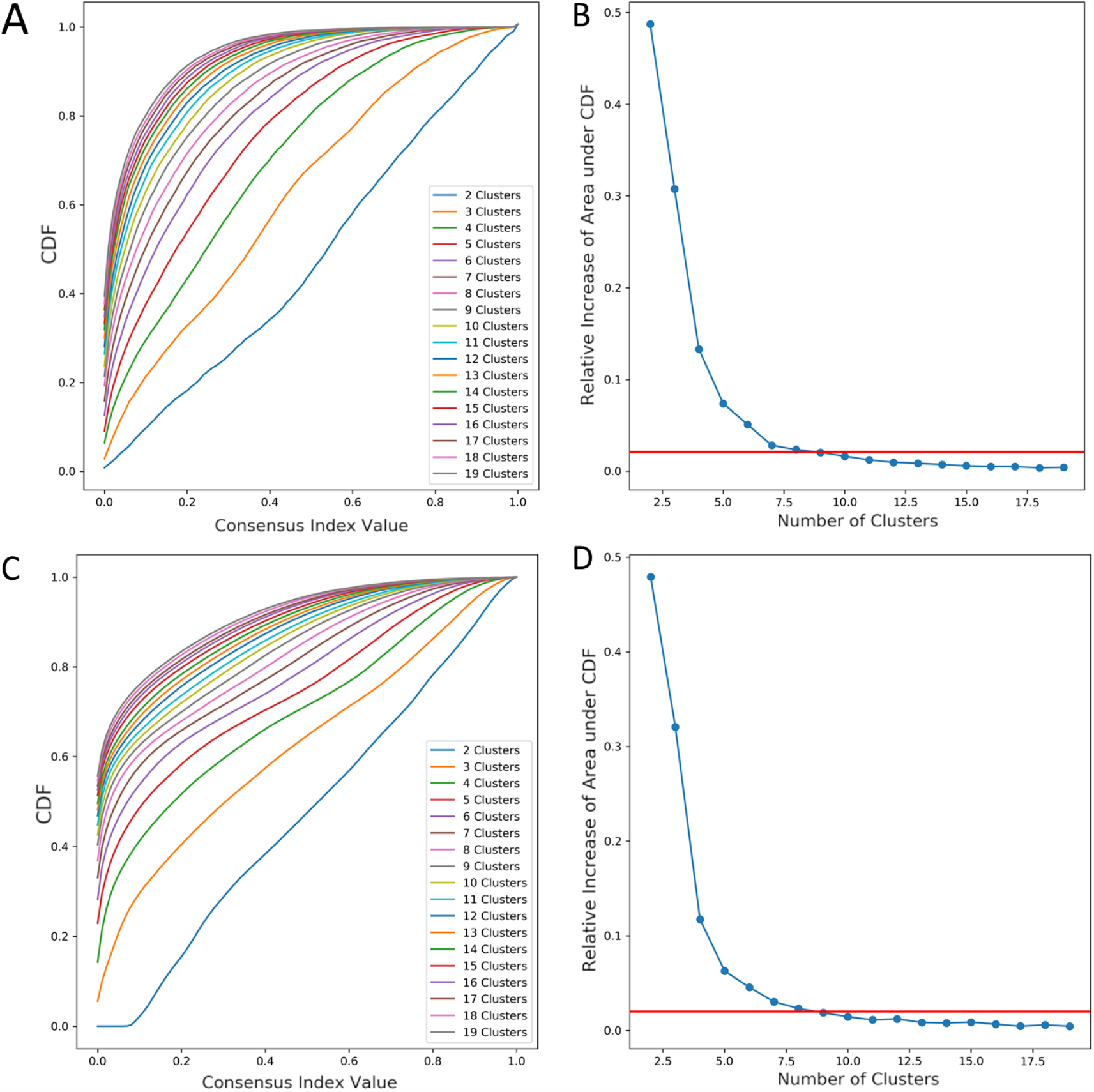
Using consensus clustering analysis for cluster number determination for the IBD (PRISM) dataset. (A) The cumulative distribution functions (CDF) for varying cluster numbers and (B) change in area under the CDF is shown for clustering on the microbial features. (C) The cumulative distribution functions (CDF) for varying cluster numbers, and (D) change in area under the CDF is shown for clustering on the metabolic features.

## Notes

### Competing Interest Statement

The authors have declared no competing interest.

